# Superior target genes and pathways for RNAi mediated pest control revealed by genome wide analysis in the red flour beetle *Tribolium castaneum*

**DOI:** 10.1101/2024.01.24.577003

**Authors:** Benjamin Buer, Jürgen Dönitz, Martin Milner, Sonja Mehlhorn, Claudia Hinners, Janna Siemanowski-Hrach, Julia K. Ulrich, Daniela Großmann, Doga Cedden, Ralf Nauen, Sven Geibel, Gregor Bucher

**Author notes:** corresponding authors Author for correspondence: Gregor Bucher. contributed equally (Benjamin Buer, Jürgen Dönitz, Martin Milner).

## Abstract

An increasing human population, the emergence of resistances against pesticides and their potential impact on the environment call for the development of new eco-friendly pest control strategies. RNA interference (RNAi) based pesticides have emerged as new option with the first products entering the market. Essentially, double stranded RNAs targeting essential genes of pests are either expressed in the plants or sprayed on their surface. Upon feeding, pests mount an RNAi response and die. However, it has remained unclear, whether RNAi based insecticides should target the same pathways as classic pesticides or whether the different mode of action would favor other processes. Moreover, there is no consensus on the best genes to be targeted. We performed a genome-wide screen in the red flour beetle to identify 905 RNAi target genes. Based on a validation screen and clustering, we identified the 192 most effective target genes in that species. The transfer to oral application in other beetle pests revealed a list of 34 superior target genes, which are an excellent starting point for application in other pests. GO and KEGG analyses of our genome wide dataset revealed that genes with high efficacy belonged mainly to basic cellular processes such as gene expression and protein homeostasis – processes not targeted by classic insecticides. In summary, our work revealed the best target genes and target processes for RNAi based pest control and we propose a procedure to transfer our short list of superior target genes to other pests.

## Background

The human population continues to grow and will reach over 9 billion by 2050 resulting in a growing demand in food supply. Together with a changing regulatory landscape and the threat of new resistances, novel crop protection solutions are required that are efficacious, durable, eco-friendly and safe to non-target organisms. RNA interference (RNAi) based solutions promise to offer such sustainable solutions with a different mode of action. However, despite many years of research there is no consensus, which genes or pathways are the best targets for RNAi mediated pest control.

RNAi is a naturally occurring defense mechanism against viruses and transposons, initially discovered in plants and in *Caenorhabditis elegans*, that has been intensively studied in various organisms. Essentially, introduction of dsRNA leads to a cellular response that destroys transcripts with sequence complementarity [1,2]. RNAi has become a valuable tool for gene function studies in many arthropods and it has been tested as new tool for species-specific and eco-friendly pest control. For instance, introduction of dsRNA targeting essential insect genes into plants has resulted in protection against pests such as the Western corn rootworm (WCR) *Diabrotica virgifera* (SmartStax™ PRO), the Cotton bollworm *Helicoverpa armigera* [3–5], and to resistance of papaya to the Papaya Ringspot Virus [6]. Many variations and formulations are being tested to increase efficacy. For instance, expression of dsRNAs in plastids led to increased lethality in the Colorado potato beetle [7]. As an alternative to *in planta* produced dsRNAs, sprayed application of dsRNA molecules onto plant surfaces has been successfully tested. The original challenge of producing dsRNA in sufficient quantity for topical applications has been solved with the advances in dsRNA production systems reducing the cost to less than US$ 0.5/g where around 0.3-4.9 g/ha are required in the field [8]. Thus, RNAi can be used for broadacre and horticultural crops through foliar spray applications. Indeed, first sprayable RNAi products are entering the market targeting Colorado potato beetle (*Leptinotarsa decemlineata*) [9] and guidelines on biosafety have been formulated [10].

Notably, the efficacy of RNAi after oral uptake widely differs across species. An efficient response has been demonstrated for several coleopteran species e.g. [4,7,11]. In *D. virgifera virgifera* and the red flour beetle *Tribolium castaneum*, targeting essential genes by RNAi results in death within a week or so [12,13] while feeding activity was reduced already after 5 days in the mustard beetle *Phaedon cochleariae* [11]. Translation to a number of other insect species including important lepidopteran pests, however, failed or led to inconsistent results e.g. [14,15]. Limitations are thought to be related to uptake and metabolism of dsRNA rather than to a general malfunction of the RNAi mechanism [16]. It has been speculated that a strong RNAi response is largely governed by efficient cellular uptake of dsRNA [17–20] and is prohibited by a high level of dsRNA degrading enzymes in the midgut and/or the hemolymph [21–26]. To overcome these challenges, new technologies for improvement of dsRNA delivery and stability are being explored [27–29].

One of the key parameters for efficacy of RNAi mediated pest control is the choice of the target genes. So far, that choice has been inspired mainly by the results from few seminal studies, which had tested a limited number of genes e.g. [4], were inspired by classic insecticidal targets or were derived from physiological knowledge e.g. [5]. However, this knowledge-based approach has been limited by the fact that most relevant parameters are unknown such as the stability of the respective protein, the dynamics of expression, and potential compensation by related proteins or regulatory mechanisms. Further, it has remained unclear whether - considering the different mode of action of RNAi - one should target different biological processes compared to the pathways known from classic chemical insecticides. Much of this uncertainty can be overcome by unbiased large-scale screening. Indeed, a previous RNAi screen in the red flour beetle *Tribolium castaneum* revealed novel target genes, which showed higher efficacy compared to previously used target genes [13,30] and they were successfully tested in other species e.g. [15,31]. Further, it identified the proteasome as an unexpected target process [13] and indeed, a proteasome component is now being used for the first sprayable dsRNA application on the market [9]. Large scale screens in other organisms had revealed essential gene sets but these screens were either not based on environmental RNAi (fly *Drosophila melanogaster*) or were performed in a species that is evolutionary quite distant to insects (nematode *Caenorhabditis elegans)* [32,33].

However, to comprehensively analyze the best target genes and processes for RNAi mediated pest control, a genome wide analysis in an appropriate laboratory model system was required. Among insects, the red flour beetle *T. castaneum* is one of the most highly developed genetic model systems and shows a strong and systemic RNAi [17,19,34,35]. Further, it is a representative of Coleoptera, which is a clade containing a number of economically relevant pests that are amenable to oral dsRNA delivery. Importantly, *T. castaneum* had been established for genome-wide screening before [13,30,36]. Some reports indicated that RNAi by feeding works in *T. castaneum* but we and others had no success in that respect [15].

To gain a genome wide view on target genes and processes, we first performed a primary RNAi screen for lethality of approximately 10,000 genes, which together with the previous screen performed by Ulrich et al. adds to an almost genome-wide coverage of 15,530 genes. Based on this primary screen, we defined a list of 905 *target genes* (top 5.8 %). GO and KEGG analyses revealed that some target processes were reminiscent of classic insecticide targets. However, most *target genes* belonged to basic cellular processes such as protein homeostasis, which are no typical targets for classic insecticides. Dose response validation of 807 of the *very good targets* and subsequent cluster analysis revealed 192 *most effective target genes* in the red flour beetle after injection of dsRNA. We then tested a subset of 66 genes by oral feeding in another beetle pest species (mustard beetle) and we found half of them to be highly active, leading to a list of 34 *superior target genes*.

Finally, we show that these *superior target genes* were very well transferable to another pest species (Colorado potato beetle). In summary, we reveal that RNAi mediated pest control should target biological processes different from chemical pesticides and we provide a list of genes, which represents an excellent starting point for identification of efficient RNAi target genes in other species.

## Results

### A very high-throughput organismal RNAi screen to detect essential genes

The realization of a genome-wide RNAi screen requires a robust experimental system. We chose the red flour beetle *T. castaneum* because this species is easy to keep in large amounts, produces offspring all year round, is easy to inject, has a robust systemic RNAi response [19,34,35] and has been established for large scale RNAi screening [12,30,36]. We needed to balance the requirements of an efficient high-throughput procedure with the aim of gathering detailed information on the dynamics at different dsRNA concentrations. To meet both needs, we opted for a two-phase screening strategy where in the primary high-throughput screen we tested all genes using one concentration of dsRNA assessing lethality at only one point in time. In the subsequent validation screen (described in the next chapter), a selection of the *target genes* identified in the primary screen were tested using different concentrations of dsRNA and scoring lethality several times after dsRNA injection (see Fig. 1A for an overview). In the primary screen, dsRNAs at a concentration of 1 µg/µl were injected into 10 larvae per gene (stages L5 or L6). The first 5,337 genes had previously been screened as part of the iBeetle screen (phase I) [30] and most of them had already been analyzed for potency as target genes for RNAi mediated pest control [13]. The main aim of the iBeetle screen had been the detection of developmental phenotypes where lethality was checked as part of an extensive morphological analysis of the injected animals at day 11 post injection [30].

**Figure 1.**
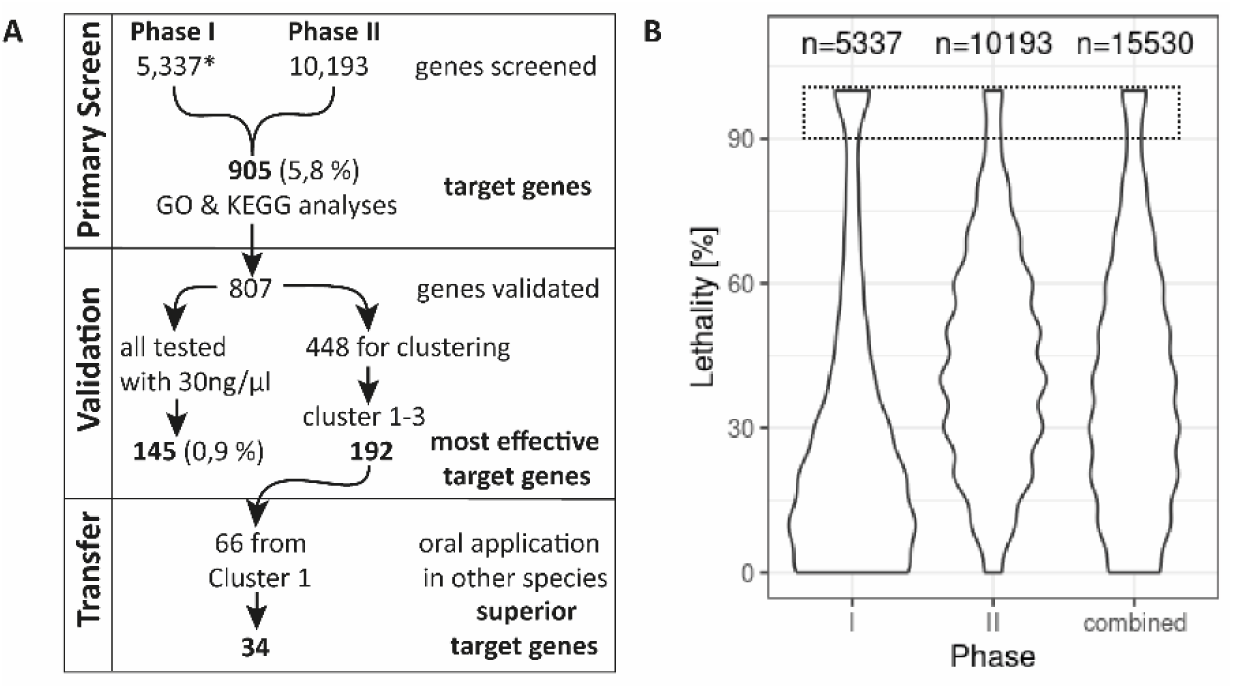
Overview and controls. A) Flowchart with the different screening phases and analyses performed in this publication. *: genes analyzed by Ulrich et al. 2015 B) Distribution of the lethality observed in phase I and phase II of the screen and for the entire dataset. The width of the shapes reflects the portion of the experiments that led to a given percentage of dead animals (n= 10 injected larvae) 11 days after injection (phase I) or 7 days after injection (phase II). Datasets with 9 or 10 dead (90 % or 100 % lethality; comprised in dotted box) were selected for the validation screen. Phase I: n = 5,337 genes; Phase II: n = 10,193 genes; total n = 15,530 genes

With this work, we present the results of phase II of the primary screen, where we scored 10,193 genes for lethality after dsRNA injection using the same concentration and the same larval stages as in phase I. However, in phase II, based on previous experience we analyzed lethality already seven days after injection to streamline the procedure. The best RNAi target genes reliably induce death within a week in *T. castaneum* such that we did not expect to miss important target genes [13]. During phase II of the primary screen, we did not score for morphological phenotypes, which allowed us to increase the throughput several-fold (see schedule and experimental details in Supporting Figure 1). While phase I had a throughput of 25 genes per week per screener (including controls and extensive phenotypic annotations [30]), we reached a throughput of 156 genes per week per screener in phase II. This increase is mainly due to the simplified readout where only the number of surviving animals was scored without preparation and morphological analyses. Based on our experience with several RNAi screens of different complexity [13,30,36], we think that this throughput is close to the upper limit for a large scale organismic RNAi screen in *T. castaneum* and probably for most if not all insects.

The distribution of the lethality found for each gene differed somewhat between phases of the screen (Fig. 1B and Fig. S2) where in phase II a lower portion of genes showed 90 % lethality or higher (compare width of distributions within the box with broken outline in Fig. 1B). This might reflect the fact that phase I of the screen was biased towards more conserved and highly expressed genes, leading to an enrichment of basic cell biological processes. On the other hand, an increased overall level of lethality was observed for phase II (Fig. 1B; see also Fig. S2). We assign the latter shift towards higher lethality in phase II to an increased technical background lethality due to stock keeping issues that we observed for some time during the screen. Of note, this technical issue did not compromise our work because subsequent selection of target genes was based on an experimentally independent validation screen (see below).

### Selection of genes for the validation screen

The primary screen provided a genome-wide insight into the characteristics of *target genes* for RNAi mediated pest control. From this set of genes, we wanted to identify the *most effective target genes*, defined as those that lead to lethality most rapidly with minimal dsRNA exposure. For this validation screen, we chose most genes that in the primary screen had shown 100 % lethality and included many genes with 90 % lethality (see Fig. S2 for more details on the selection). Specifically, from the 623 and 282 genes that had shown 100 % or 90 % lethality in the primary screen, respectively, 607 and 200 genes were included. In summary, 807 of these 905 genes (further referred to as “*target genes”)* were included in the validation screen, i.e. they were tested in independent experiments using lower concentrations of dsRNA and monitoring the lethality over time. In addition, 16 genes with lower lethality were tested to check for consistency with the results of the primary screen.

### Validation screen to detect the most efficient RNAi target genes

All 807 genes included in the validation screen were injected with a dsRNA concentration of 30 ng/µl and subsets were treated in addition with 3 ng/µl or 300 ng/µl dsRNA. The lethality distributions revealed that 300 ng/µl closely reflected the results found in the primary screen (1 µg/µl) in that a high degree of lethality of almost all tested genes was observed after 6-8 days (Fig. 2A, right panel). At 30 ng/µl concentration of injected dsRNA, the knock-down led to lethality in many but not all genes (Fig. 2A middle panel). The lowest concentration (3 ng/µl) induced lethality in a yet smaller portion of genes (Fig. 2A left panel). We concluded, that the concentration of 300 ng/µl was too high to be a stringent selection criterion while 3 ng/µl was close to the lower limit of dsRNA concentration that can induce a lethal effect in *T. castaneum* by injection.

**Figure 2:**
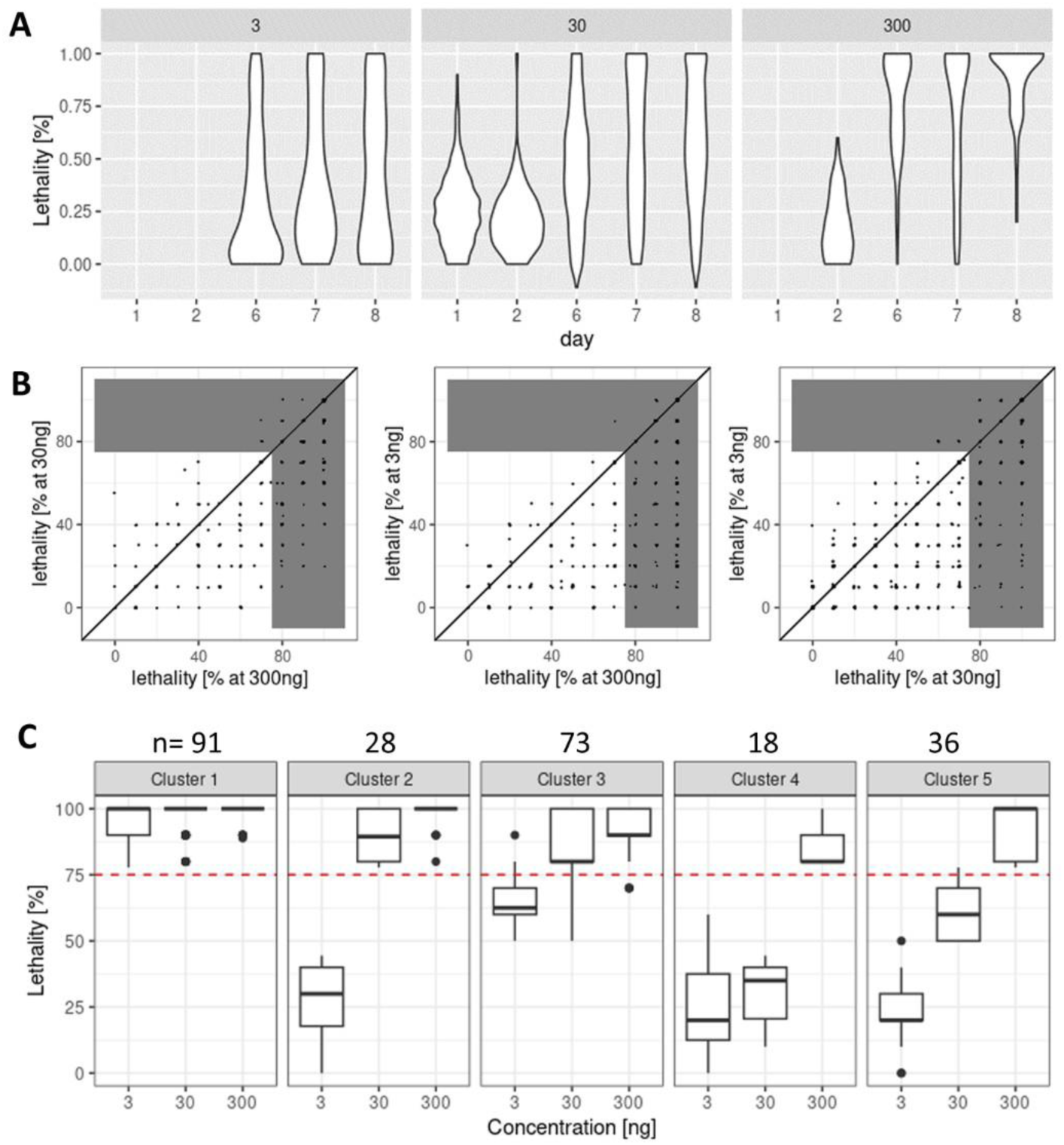
Distribution and correlation of lethality at different concentrations found in the validation screen. (A) The lethality distribution found in the validation screen is depicted along the time axis (in days after injection; see Supporting Figure 3 for separate documentation of the results of different validation screen phases). As expected, the effect is dose and time dependent. Notably, the distributions found with the lowest concentration (left panel) differ less along the time axis compared to higher concentrations. (B) Correlation of lethality between different concentrations of dsRNA targeting the same gene at 7 or 8 days post injection. Each dot reflects the lethality induced by a given dsRNA injected at the different concentrations (see axes). Lethality equal or greater than 75 % is shaded in grey. As expected, the lethality increased with the concentration. The correlation of the two high concentrations (left panel) was high while the lowest concentration (3ng/µl) failed to induce lethality for a large number of genes, which did have strong effects at 30 or 300 ng/µl (see dots in right bottom part of grey area in the middle and right panels). Nevertheless, a number of genes induced lethality even at the lowest concentration (top right grey area). (C) Cluster analysis of a subset of the genes tested in the validation screen according to their level of lethality. Clusters 1-3 were used to define the *most effective target genes* used for subsequent analyses. See text for further details.

Analysis of the lethality induced by different concentrations of the same dsRNA essentially confirmed these results (Fig. 2B). While the lethality at the higher concentrations correlated (Fig. 2B, left panel), many dsRNA that induced lethality at higher concentrations failed to do so at 3 ng/µl (Fig. 2B, middle and right panels).

### Defining the most effective RNAi target genes

Based on our data, we considered two ways to define the *most effective target genes* from the validation-screen. In the first approach, we used only the results from experiments with 30 ng/µl concentration because this concentration had been used for all genes in the validation screen. From all these genes, 145 showed a lethality of 100 % (0,9 % of all genes of the primary screen) (see Supporting Table 3 for gene IDs). While this approach included all genes from the validation screen, it did not take into account dose dependent responses and was therefore not used for follow-up experiments. In an alternative approach, we performed cluster analysis on the subset of 246 genes that had been validated with 3 ng/µl and that showed a lethality ≥ 75% at the 30 ng/µl concentration. Unsupervised K-means clustering was performed based on the percent lethality at different concentrations at day 7 or 8 after injection with dsRNA. We obtained five distinct clusters with Cluster 1 comprising the 91 most potent target genes with high lethality at all concentrations (Fig. 2C). Similarly, Cluster 3 (73 genes) was comprised of highly potent targets with some decline of efficiency at 3 ng/µl dsRNA injection. Cluster 2 (28 genes) showed more pronounced cutoff concentrations. Cluster 5 (36 genes) shows the clearest dose response. Cluster 4 (18 genes) shows lowest efficacy. See Supplementary Table 4 for the gene IDs of these clusters, Supplementary Table 5 for respective GO-term and KEGG analyses and the top 15 GO terms of Cluster 1.

Taken together, our genome wide screen by injection in the red flour beetle revealed a list of 905 *target genes*, which represented an excellent basis for understanding biological processes targeted by these genes. The validation screen and cluster analysis led to the identification of 192 *most effective target genes* derived from clusters 1-3. This set of genes represented an excellent starting point for transferring our findings to relevant pests by administering dsRNA by feeding.

### The majority of the target genes are part of basic cellular processes

To reveal the biological processes and pathways, which should be targeted in RNAi mediated pest control, we asked, which functions and pathways were enriched in the *very good target gene* set defined in our primary screen (905 genes representing 5.8 % of all screened genes) compared with the set of genes with a mortality of 50 % or less. The genes were annotated by similarity and functional domains using blast2go and mapped to KEGG pathways [37,38]. Gene Ontology (GO) term enrichment analysis resulted in 393 enriched GO terms represented by at least 5 genes with p-values ≤ 0.01. The top GO terms according to their p-value are listed in Table 1 (see Supplementary Table 1 for all enriched GO terms; the IDs of the genes contributing to the top 15 GO annotations are found in Supplementary Table 2). The top GO terms in the domain “biological process” reflected predominantly basic cellular processes involved in gene expression such as translation, transcription and RNA metabolism (Table 1). Two terms were related to development but most of the underlying genes are involved in basic cellular processes that indirectly influence developmental pattern formation (see Supplementary Table 2 for gene IDs). Notable exceptions are the nuclear hormone receptor Ftz-F1 (TC002550) and the Glycogen synthase kinase-3 beta (TC032822), which have direct functions in pattern formation in addition to metabolic processes [39]. The most enriched GO terms of the domains “molecular function” and “cellular component” reflected the above-mentioned findings. The enriched cellular component “plasmodesma”, which actually is a plant-specific structure, may reveal unspecific functional annotation and hint towards enrichment. of the term “cell-cell junction” (GO:0005911) which is a parent GO term of both, plasmodesma and cell-cell structures from animals.

**Table 1:**
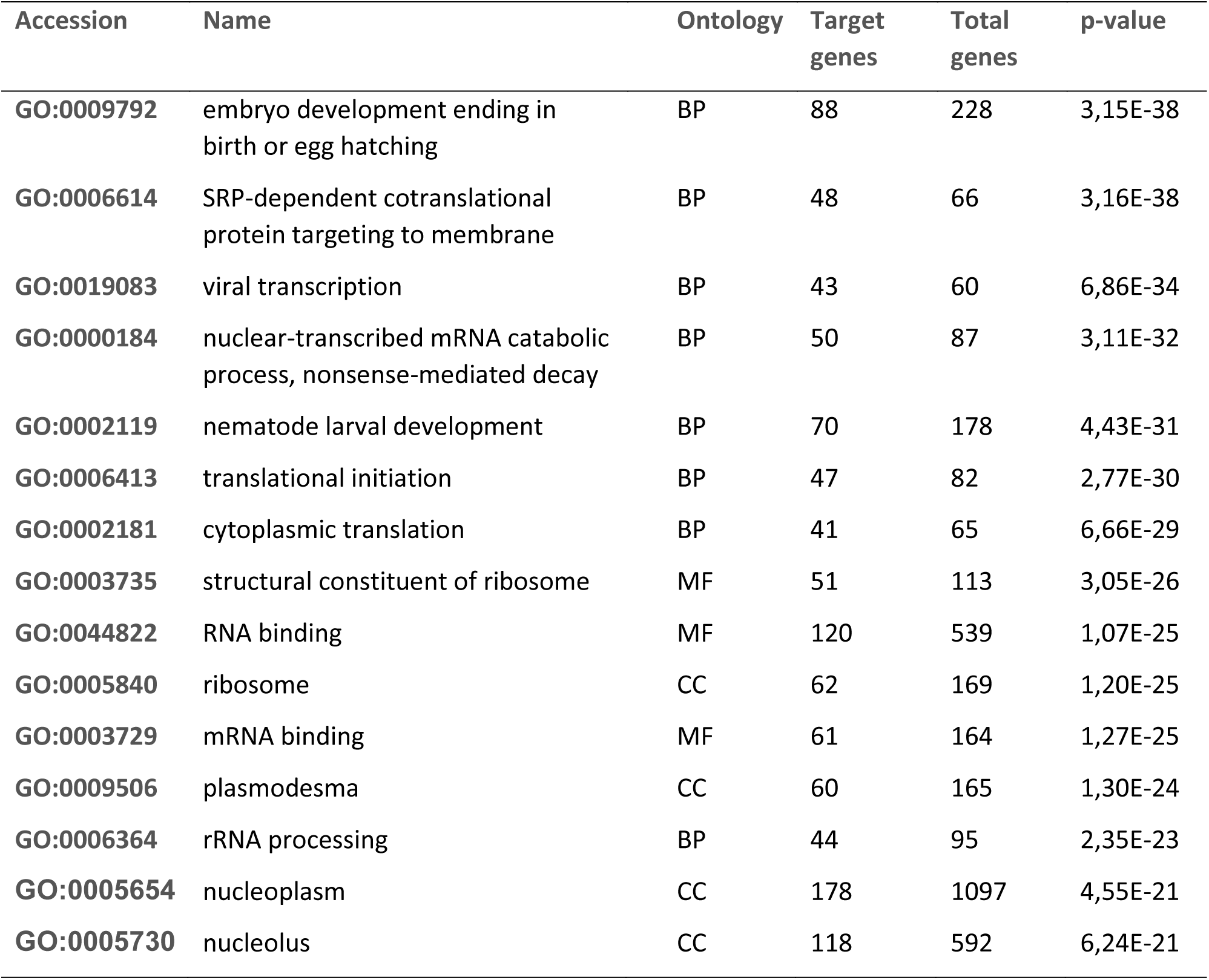
Gene ontology enrichment of the set of *very good RNAi target genes*. The 905 genes with mortality larger or equal 90 % in the primary screen were compared to all genes with mortality less or equal 50 % using hypergeometric distribution. Top 15 enriched GO terms are shown. See Supplementary Table 1 for all GO terms and Supplementary Table 2 for respective gene IDs.

| Accession | Name | Ontology | Target genes | Total genes | p-value |
| --- | --- | --- | --- | --- | --- |
| GO:0009792 | embryo development ending in birth or egg hatching | BP | 88 | 228 | 3,15E-38 |
| GO:0006614 | SRP-dependent cotranslational protein targeting to membrane | BP | 48 | 66 | 3,16E-38 |
| GO:0019083 | viral transcription | BP | 43 | 60 | 6,86E-34 |
| GO:0000184 | nuclear-transcribed mRNA catabolic process, nonsense-mediated decay | BP | 50 | 87 | 3,11E-32 |
| GO:0002119 | nematode larval development | BP | 70 | 178 | 4,43E-31 |
| GO:0006413 | translational initiation | BP | 47 | 82 | 2,77E-30 |
| GO:0002181 | cytoplasmic translation | BP | 41 | 65 | 6,66E-29 |
| GO:0003735 | structural constituent of ribosome | MF | 51 | 113 | 3,05E-26 |
| GO:0044822 | RNA binding | MF | 120 | 539 | 1,07E-25 |
| GO:0005840 | ribosome | CC | 62 | 169 | 1,20E-25 |
| GO:0003729 | mRNA binding | MF | 61 | 164 | 1,27E-25 |
| GO:0009506 | plasmodesma | CC | 60 | 165 | 1,30E-24 |
| GO:0006364 | rRNA processing | BP | 44 | 95 | 2,35E-23 |
| GO:0005654 | nucleoplasm | CC | 178 | 1097 | 4,55E-21 |
| GO:0005730 | nucleolus | CC | 118 | 592 | 6,24E-21 |

We asked, whether signaling pathways would be good targets due to their involvement in many biological processes. We found the NIK/NF-kappa B, TNF, Wnt and MAPK signaling pathways to be significantly enriched in our set of candidate target genes but they were not among the top 15 (see Supplementary Table 2). Further GO terms associated with molecular functions comprised protein binding and cell-cell adhesion as well as endocytosis and proton-transporting ATPase activity. In line with the latter term, V-ATPase had been introduced as a potent RNAi target gene before [4] and has been successfully used by others since. The previously described enrichment of proteasome components in RNAi target genes [13] was found in this genome wide dataset as well, albeit not with the highest scores. Other enriched GO terms in the cellular component domain included key cellular complexes such as the proteasome and ribosome and general compartments such as the vacuolar membrane, cytoplasm and nucleoplasm and specialized structures such as the myelin sheath and exosome.

To map and visualize enriched GO terms and their functional interconnections, we created networks using REVIGO and GO slim annotations (Supplementary Figures 4-6) [40]. With respect to biological process, the network consists of two major subnetworks: One reflecting regulatory processes (top part in Supplementary Fig. 4) and one reflecting transcription, translation and related decay processes (center part in Supplementary Fig. 4). With respect to the category “cellular component”, only one network was found that included mainly translation and protein decay (Supporting Fig. 5).

In summary, our GO term analysis revealed that the set of *target genes* identified from a genome wide screen was highly enriched in basic cellular processes with translation, protein homeostasis, transcription and RNA biology being among the top processes. Our analysis shows that genes in biosynthetic processes are more abundant among the most efficient target genes than genes for structural components, which were not much enriched in our dataset.

In order to complement the functional enrichment with pathway information, our collection of *target genes* was assigned to KEGG pathways. Essentially, this analysis yielded pathways similar to those found by the GO term enrichment, e.g. ribosome, proteasome and spliceosome as top annotations and a number of additional processes related to protein and RNA biology (Table 2).

**Table 2:** KEGG enrichment of the set of *target genes*. The 905 genes with mortality larger or equal 90% in the primary screen were analyzed for annotation in KEGG pathways.

| Accession | Name | Lethal genes | Total genes | p-value | Lethal genes (%) |
| --- | --- | --- | --- | --- | --- |
| ko03010 | Ribosome | 43 | 69 | 1,72E-18 | 62.3 |
| ko00190 | Oxidative phosphorylation | 30 | 58 | 2,50E-10 | 51.7 |
| ko03050 | Proteasome | 22 | 35 | 4,50E-10 | 62.9 |
| ko03040 | Spliceosome | 35 | 82 | 6,01E-09 | 42.7 |
| ko04145 | Phagosome | 24 | 52 | 2,81E-07 | 46.2 |
| ko04721 | Synaptic vesicle cycle | 16 | 30 | 2,69E-06 | 53.3 |
| ko03060 | Protein export | 11 | 16 | 3,54E-06 | 68.8 |
| ko03013 | RNA transport | 27 | 73 | 1,00E-05 | 37.0 |
| ko00970 | Aminoacyl-tRNA biosynthesis | 12 | 25 | 0,00018991 | 48.0 |
| ko03020 | RNA polymerase | 9 | 16 | 0,00027484 | 56.3 |
| ko03022 | Basal transcription factors | 11 | 25 | 0,00089401 | 44.0 |
| ko03015 | mRNA surveillance pathway | 16 | 46 | 0,00146617 | 34.8 |
| ko03008 | Ribosome biogenesis in eukaryotes | 14 | 44 | 0,00716605 | 31.8 |
| ko04623 | Cytosolic DNA-sensing pathway | 7 | 16 | 0,0083228 | 43.8 |
| ko04714 | Thermogenesis | 26 | 102 | 0,00923878 | 25.5 |

Interestingly, in this analysis the proteasome was recovered as one of the three top pathways reflecting our previous findings [13]. Notably, the KEGG pathway ‘oxidative phosphorylation’ had a high score indicating that energy metabolism may be a good target for RNAi mediated pest control as well.

### Transfer to oral feeding in other pest species reveals 34 *superior target genes*

We tested some of the *most effective target genes* in other species for two reasons. First, our genome-wide screen was based on injection of dsRNA, which probably has different characteristics compared to oral application. Oral application of dsRNA did not work for *T. castaneum* in our hands despite positive reports from others [41–44]. Second, while some RNAi target genes had successfully been transferred to other species, limits of transferability had been observed as well. For instance, some of the top target genes from our previous study [13] had been tested in other species where some but not all turned out to be effective [15]. That indicated that the transferability of a given gene may be dependent on the species and we could not identify a clear pattern of effectiveness for these target genes in that survey [15]. Therefore, we tested for the transferability of *superior target genes* to different species and changing the delivery mode to oral application.

We used the mustard beetle *Phaedon cochleariae*, which is a well-described pest that had previously been established for RNAi screening [11]. We randomly chose 88 genes from our *most effective target genes* (i.e. clusters 1 – 3), determined their orthologs in *P. cochleariae* and tested respective dsRNAs by oral delivery. We used a concentration corresponding to 30 g/ha or 100 g/ha and checked for effects after 10 days maximum. We tested 66 sequences from cluster 1 and found that 34 (52 %) showed increased lethality by more than 50 % in comparison to controls (Figure 3A). For cluster 2 (17 genes) the rate declined to 18 % while for cluster 3 (5 genes) we found 40 %. These results underscored the variability of the transfer across species and/or delivery modes. Based on this experiment we defined the 34 successfully transferred genes from Cluster 1 to be our final selection of *superior target genes*. This set of genes represents a manageable number to be tested in other pest species yet may be large enough to accommodate for species-specific variability. The gene IDs of the 34 *superior target genes* are given in Table 3. Note that additional similarly effective target genes might be present, e.g. in the 26 non-tested genes from clusters 1-3 or in the set of most efficient target genes that we defined by the alternative approach (see above).

**Figure 3.**
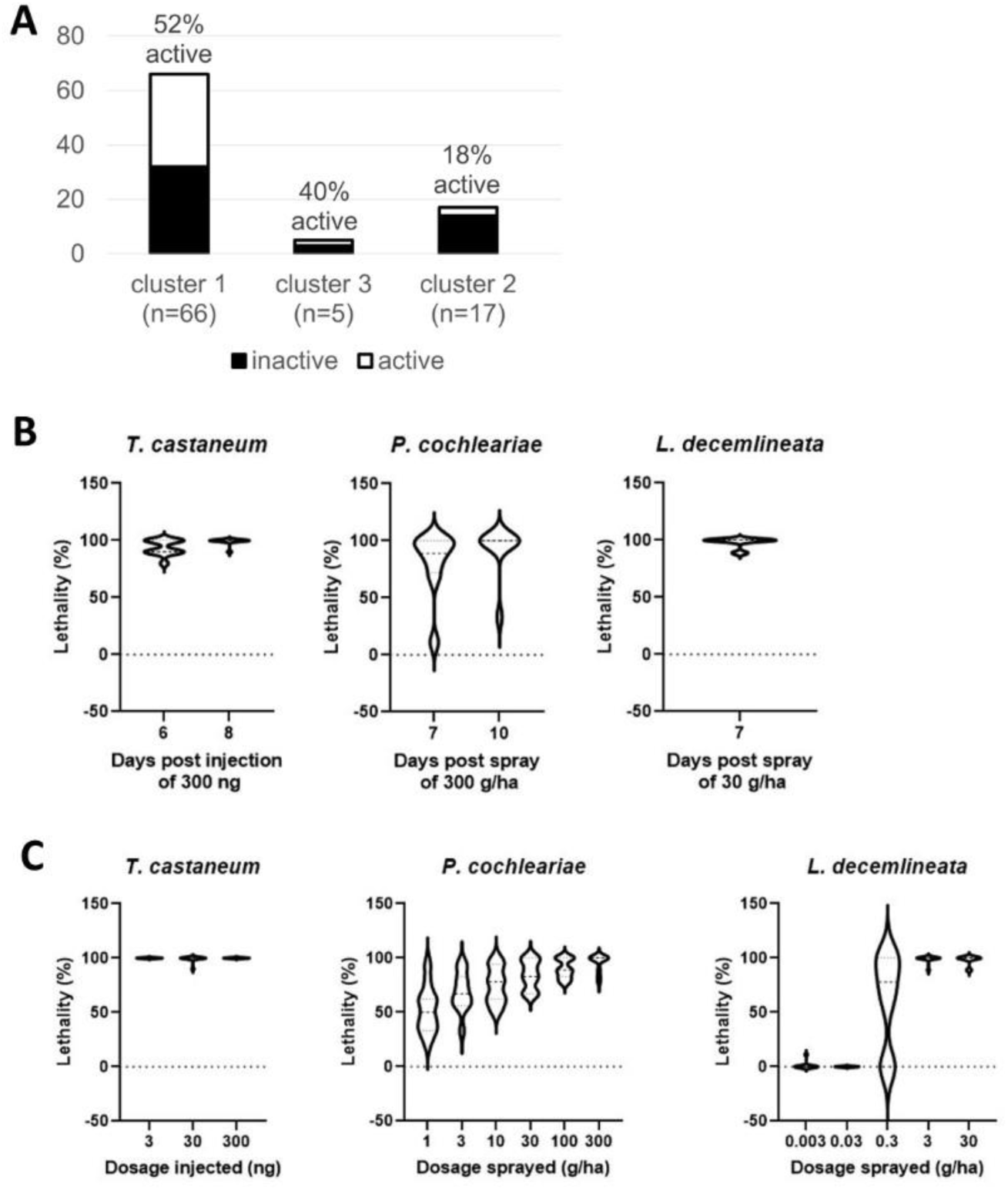
Transfer of most effective target genes to other species. A) From 66 genes of cluster 1, we found 52% to be transferable to P. cochleariae by oral feeding. The transferability for the other clusters was lower. We defined the 34 genes transferred from Cluster 1 to represent our superior target gene set. B) Time to mortality induced by 11 selected genes that were active in T. castaneum, P. cochleariae and L. decemlineata. Lethality was found within 7-10 days in all three species. C) These 11 genes were sensitive to small amounts of dsRNA but large species-specific differences of sensitivity were observed.

**Table 3.** Superior target genes recommended for transfer to other pest species. These 34 genes were identified as most efficient target genes in T. castaneum by injection (i.e. they are part of Cluster 1) and were successfully transferred to C. cochleariae by feeding. The 11 genes shaded in grey were additionally successfully transferred to L. decemlineata.

| Tribolium gene ID | iBeetle number | Gene name |
| --- | --- | --- |
| TC000069 | iB_00011 | Proteasome subunit beta type-1 |
| TC000614 | iB_00141 | Proteasome subunit alpha type-6 |
| TC000641 | iB_00148 | Coatomer subunit beta |
| TC002574 | iB_00404 | Signal recognition particle 54 kDa Short=SRP54 |
| TC004425 | iB_03754 | Heat shock 70 kDa cognate 3 |
| TC005185 | iB_06598 | Polyadenylate-binding 1 Short=PABP-1 Short=Poly(A)-binding 1 |
| TC005653 | iB_07516 | snakeskin |
| TC006375 | iB_04125 | 26S proteasome non-ATPase regulatory subunit 6 |
| TC006492 | iB_04154 | 26S proteasome regulatory subunit 7 |
| TC007891 | iB_01280 | 26S proteasome non-ATPase regulatory subunit 8 |
| TC007999 | iB_04411 | 26S proteasome regulatory subunit 6B |
| TC008617 | iB_01375 | Proteasome subunit beta type-4 |
| TC009191 | iB_01493 | transport Sec23A |
| TC009491 | iB_01562 | RNA-binding 15 |
| TC009675 | iB_08675 | 26S proteasome regulatory subunit 4 Short=P26s4 |
| TC009965 | iB_04734 | Ras-related Rab-1A |
| TC010003 | iB_01640 | DNA-directed RNA polymerases I, II, and III subunit RPABC1 |
| TC010318 | iB_01665 | Bifunctional 3'-phosphoadenosine 5'-phosphosulfate synthase (PAPS synthase) |
| TC010321 | iB_04808 | 26S proteasome regulatory subunit 10B |
| TC010519 | iB_01704 | ATP-binding cassette sub-family E member 1 |
| TC011058 | iB_01793 | Dynamin |
| TC011120 | iB_01807 | ROP |
| TC011182 | iB_01820 | 60S ribosomal L7a |
| TC012303 | iB_01965 | Eukaryotic translation initiation factor 3 subunit A (eIF3a) |
| TC013782 | iB_02204 | Glycine--tRNA ligase |
| TC014413 | iB_05628 | ATP-dependent RNA helicase WM6 Short=DEAD box UAP56 Short=Dmrnahel |
| TC014725 | iB_09124 | Prolactin regulatory element-binding |
| TC015014 | iB_02377 | Clathrin heavy chain |
| TC015205 | iB_08499 | Proteasome subunit beta type-3 |
| TC015539 | iB_09161 | 40S ribosomal S3a |
| TC031132 | iB_07271 | Pre-mRNA-splicing factor SYF1 |
| TC033036 | iB_02787 | 26S proteasome non-ATPase regulatory subunit 1 |
| TC034312 | iB_09459 | Splicing factor 3B subunit 1 |
| TC034766 | iB_01582 | Tubulin beta-3 chain |

To confirm transferability of the *superior target genes*, we tested a subset of 12 genes in the *L. decemlineata,* an organism that had shown excellent response to RNAi [45]. Indeed, 11 out of 12 sequences showed strong effects compared to our controls indicating a high degree of transferability (marked in grey in Table 3). Interestingly, these eleven genes led to lethality within 7-10 days in all tested species (Fig. 3C).

## Discussion

### RNAi mediated pest control can target biological processes not used by classic insecticides

With this work, we present the first genome-wide screen of the most effective target genes and pathways. Our study provides an unbiased whole genome view whereas many other studies chose to target classic insecticidal targets with RNAi. Indeed, we detected novel target pathways only some of which had been known from classic insecticidal processes such as synaptic vesicle cycle or oxidative phosphorylation. However, most *target genes* acted in basic cellular processes such as transcription, protein translation, export and degradation. Notably, these processes are very different from the modes of action of classic insecticides. A reason for this discrepancy might be that the protein domains essential for basic cellular processes are often highly conserved between insects and vertebrates such that most chemical inhibitors would not pass the biosafety measures. Moreover, RNAi might be able to target the expression of genes whose protein products are not accessible for classic insecticides due to their cellular localization or quaternary structures known for instance from the proteasome or ribosomal proteins. To avoid cross-effects on non-target organisms, RNAi can be directed to diverged sequences including UTRs, which allows for species-specific targeting of even highly conserved proteins, enabling bio-safe targeting [10,13]. Indeed, the pilot for this screen had identified the proteasome as prime target and the first sprayable application is based on a proteasome subunit [9,13]. In summary, RNAi opens essential basic cellular pathways for targeting, which have been protected from classic insecticides.

### How comprehensive was our analysis?

The primary screen (15.530 genes) covered 93,6 % of the current protein coding gene set of the *T. castaneum* genome assembly OGS3 [46] and mainly missed genes that could not be cloned from cDNA and genes that were affected by the usual loss of experiments during high throughput screens (see Supplementary Table 6 for all results of the primary screen). Therefore, our list of 905 *target genes* (top 5.8 % of the tested genes) is very comprehensive and our conclusions on GO terms and KEGG pathways provide the first and a very robust genome-wide view on that matter. Likewise, the validation screen was quite comprehensive where 807 out of 905 *target genes* were tested (89,2%).

The subsequent steps had the aim of identifying a manageable number of superior target genes for transfer to other pests rather than providing comprehensive analyses. Therefore, the cluster analysis was based only on those 443 genes, which had been validated with the lowest concentration (54,9 % of 807 genes in the validation screen). Due to this restriction, about half of the genes matching our criteria for *superior target genes* are probably missing from our list. Likewise, we tested a subset of 66 out of 91 genes from Cluster 1 (72,5 %) to define our list of *superior target genes*. This means that another dozen genes or so from Cluster 1 may show a similar efficacy when transferred to other species. Given the high transferability of our *superior target genes* to another pest species (11 out of 12; 91,7 %) we think that 34 genes are a sufficient and at the same time manageable number. If testing of the entire *superior gene* list does not result in an efficient RNAi response in an organism, the root cause is likely to lie in other reasons than the selection of the appropriate target gene.

### Comparison of our superior target genes to the target genes currently used

A number of genes had previously been used by others as targets for RNAi mediated pest control and we asked, how far these genes were comprised in our lists. Importantly, several popular target genes such as chitin synthase, acetylcholinesterase or ecdysone receptor were found in none of our lists indicating that there is room for improvement for respective applications by testing our *superior target genes*.

Three previously used genes were in our *superior target gene* list: Sec23, heat shock 70 kDa, and COPI coatomer β subunit (bold in Table 4) confirming that these previous screening efforts have identified excellent target genes. Neither of the currently registered RNAi-based products and only 5 out of 12 commonly used target genes belonged to our top performing Cluster 1 genes (see Table 4). However, another three represented different subunits of protein complexes that were targeted by at least one of our *superior target genes*: The proteasome, microtubules and the ribosome (marked in grey in Table 4). While these genes are likely quite good target genes, testing the *superior target genes* as targets would still be advisable as even a minor increase of efficacy reduces the cost for application. The most lethal eleven genes identified in a previous screen belonged to Cluster 1 while a machine learning approach to identify essential genes had revealed only one Cluster 1 target gene.

**Table 4.**
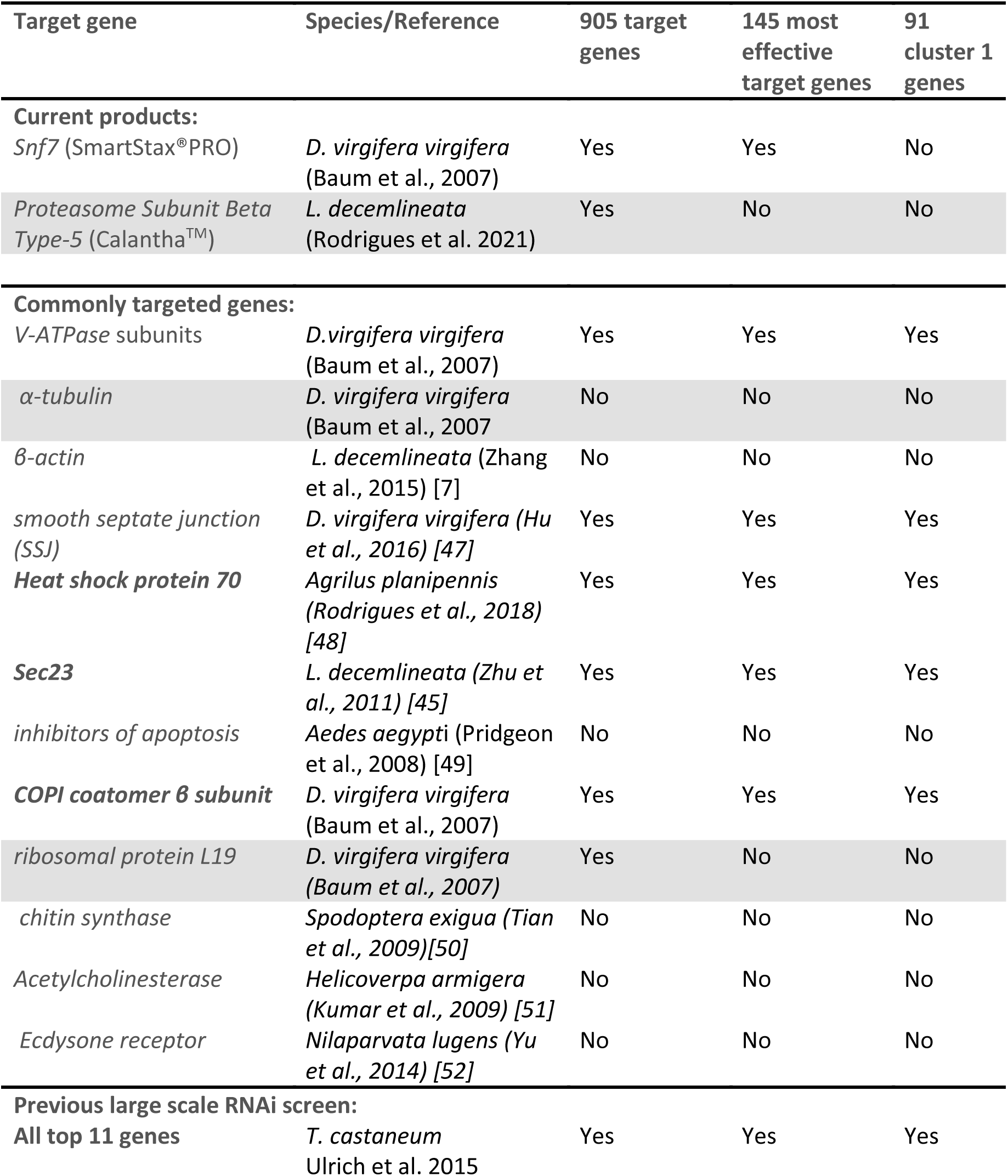
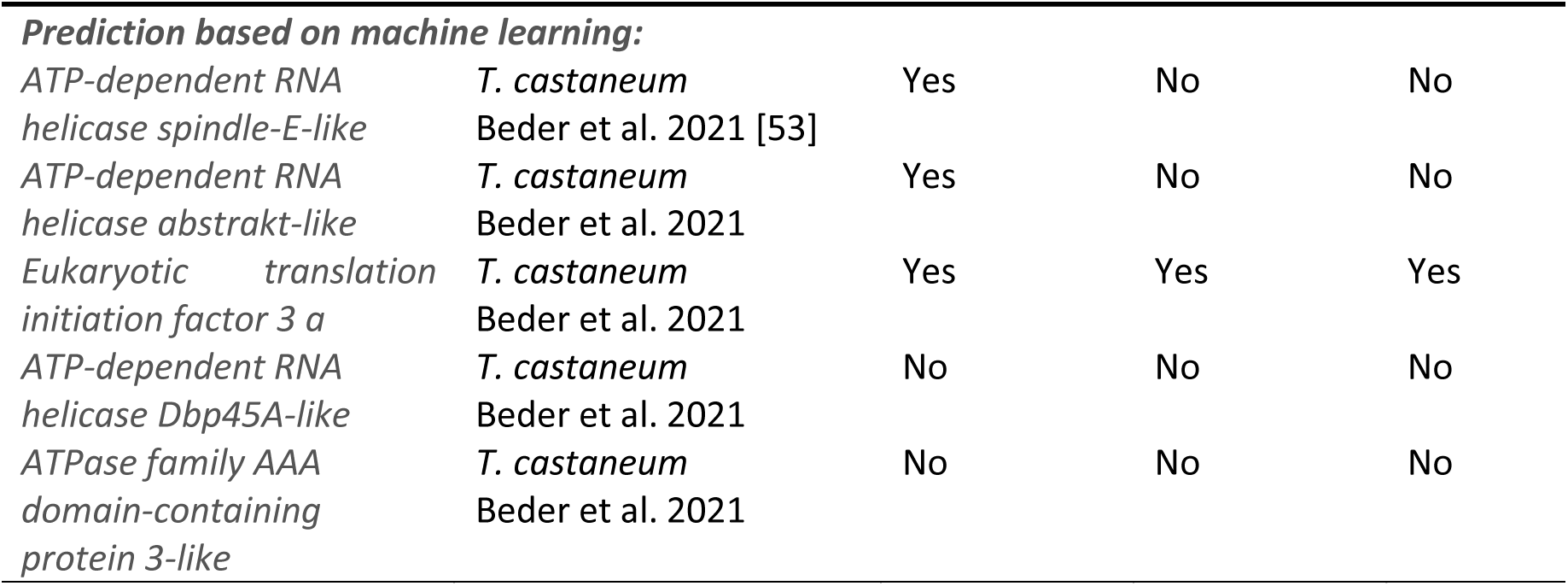
Commonly used target genes compared to our gene sets. Selected target genes were checked whether they were included in the 905 *target genes* (identified in the primary screen), the 145 *most effective target genes* (based on the validation screen based on 30ng/ul; see above) or the 91 genes from Cluster 1. Three genes previously published by others were included in our *superior target gene* list (shown in bold) and three targeted different subunits of the protein complexes targeted by *superior target genes* (shaded in grey).

In summary, our superior target genes perform better than many genes previously selected based on knowledge on protein functions - at least under the tested conditions such as species, targeted stage or selected sequence. One reason could be that the knowledge-based approach does not take into account additional parameters that influence the RNAi response. For instance, high protein stability increases the time from knock-down to biological effect; alternative pathways or paralogs may compensate for the loss of an essential protein and compensatory upregulation of expression may counteract the knock-down effect. Moreover, we do not know, in which cell types a given gene may be essential and which cells take up dsRNA efficiently. Given our lack of knowledge of most parameters for most of the genes, our unbiased large-scale screen seemed the tool of choice for the identification of the most efficient target genes. Interestingly, both current RNAi-based commercial products are targeting genes that are comprised in our *very good target gene* set (905 genes with >90 % lethality) and one of them, Snf7 (SmartStax®PRO), even belonged to the *most effective genes*. This highlights the potential for transferability from an unbiased screen in a model species to the market.

### Considerations for the identification of the most effective target genes for RNAi mediated pest control

We propose keeping the following considerations in mind when planning to identify the target genes for RNAi mediated pest control:

First, the selection of a target gene based on the knowledge of its essential function is often suboptimal because we lack the knowledge of all the other parameters influencing an efficient RNAi response such as the developmental stage, dsRNA stability etc. For instance, some excellent targets for chemical insecticides have performed poorly when targeted by RNAi.

Second, due to species-specific variation of parameters influencing the effect of RNAi, a gene with an excellent response in one species may be less effective in another. Hence, there is no such thing as *the one best target gene*. See table in Mehlhorn et al. 2021b for examples. As consequence, several putatively efficient target genes have to be tested rather than relying on one.

Third, our list of *superior target genes* is an excellent starting point for a small scale screen to identify the best targets in another species. Such an approach focuses on some of the most promising targets but it still considers the possibility of species-specific variability. Testing 34 genes will be realistic for most systems and we consider the likelihood to be rather high that at least one of them will belong to the species-specific top group. If a more comprehensive approach is needed, the remaining genes of Cluster 1 could be included or the 145 genes of the *most effective target genes* could be tested.

Fourth, careful controls and independent replicates are paramount to avoid false-positive reports on RNAi in pest control. Lethality is a very unspecific phenotype that is often elicited by a variety of technical variables such as contaminated injection needles, poor dsRNA preparations, stock keeping issues, infection status etc. Before testing RNAi for pest control with such an unspecific readout, an efficient RNAi response should first be confirmed. To that end, the use of non-lethal target genes with a clear phenotypic readout such as pigmentation genes are advisable. See Mehlhorn et al. 2021b for suggestions.

Fifth, many of the target processes are highly conserved in eukaryotes. Hence, they might be valuable targets in other economically relevant arthropods such as spider mites or even other clades of eukaryotes such as fungi.

## Conclusions

Our genome-wide approach allowed us making well-founded statements on the processes and genes that are the best targets for RNAi based pest control. We found that most of the best RNAi target genes are highly conserved genes acting in basic cellular and biosynthetic processes. Further, we provided a short list of superior target genes, which can be used as starting point for future efforts to establish this technique in other pest species. While it may seem unlikely to identify more efficient protein coding target genes, it remains elusive how well non-coding RNA targets may perform. Another future main challenge will be to increase the efficacy of the specific target sequence by rational design.

## Material and methods

### Primary screen

The screening followed the procedure extensively described in Schmitt-Engel et al. 2015 with minor modifications. In brief, pBA19 L6 or L5 instar larvae were injected and scored for lethality. dsRNA solution at a concentration 1µg/µl was injected into 10 animals per experiment (mixed males and females). Per screening day, injections for 40 different genes and controls were performed, where the first round of injection of each day represented the negative control (injection buffer). Injections were performed four days a week with the fifth day being required for stock maintenance and documentation. Using this schedule, each week 156 novel genes were injected. On day 7 and 16 after injection, the lethality of the larvae, pupae and adults, respectively was determined. See supporting Figures 1 and 5 for detailed information.

### Functional annotation of genes

Functional annotation of *Tribolium castaneum* genes was performed as described in [54]. In brief, BLAST2GO v1.3.3 [55] was used to summarize annotation of protein domain predictions from InterproScan v5.17-56.0 [56] and similarity searches against Uniprot KB using NCBI-BlastP v2.2.27 [57].

Functional and pathway enrichment analysis was performed using hypergeometric distribution with R package goseq v1.28.0 [58] with default parameters. For enrichment of pathways, annotation from KEGG database was used [38].

For visualization and clustering of enriched GO terms, REVIGO v1.8.1 [40] was used with default parameters. Clustering of GO terms was performed at a cutoff value of 0.9.

### Clustering

Clustering of lethal genes was performed using kmeans clustering with R v3.6.2. The number (n=5) of clusters was defined using the elbow method. A number of genes had not been screened with 300 ng/µl concentration. In order to include them in the analysis, we replaced these values with the values from the 30 ng/µl concentration. Based on our previous distribution and correlation analyses (Fig. 2A,B) this should not introduce a concerning bias that would interfere with our aims.

## Supporting information

GO terms and KEGG pathways enriched in very good target genes

Gene lists underlying the top 15 GO terms

Most effective target genes based on lethality at 30ng_per_ul

Most effective target genes based on clustering

GO and KEGG enrichment analysis of clusters 1-5

Lethality results of the primary screen for all genes

## Supporting information

Supporting file 1: Supporting text and figures 1 6

Supplementary Table 1: GO terms and KEGG pathways enriched in very good target genes

Supplementary Table 2: Gene lists underlying the top 15 GO terms

Supplementary Table 3: *Most effective target genes* based on lethality at 30ng_per_ul

Supplementary Table 4: *Most effective target genes* based on clustering

Supplementary Table 5: GO and KEGG enrichment analysis of clusters 1-5

Supplementary Table 6: Lethality results of the primary screen for all genes

## Acknowledgements

This work was supported by the DFG research unit FOR1234 *iBeetle* via the grants BU1443/6-1 and 6-2. We thank Elke Küster for help with the screen.

## References

1. Fire A, Xu S, Montgomery MK, Kostas SA, Driver SE, Mello CC. Potent and specific genetic interference by double-stranded RNA in Caenorhabditis elegans. Nature. 1998;391:806–11.

2. Meister G, Tuschl T. Mechanisms of gene silencing by double-stranded RNA. Nature. 2004;431:343–9.

3. Bachman PM, Bolognesi R, Moar WJ, Mueller GM, Paradise MS, Ramaseshadri P, et al. Characterization of the spectrum of insecticidal activity of a double-stranded RNA with targeted activity against Western Corn Rootworm (Diabrotica virgifera virgifera LeConte). Transgenic Res. 2013;22:1207–22.

4. Baum JA, Bogaert T, Clinton W, Heck GR, Feldmann P, Ilagan O, et al. Control of coleopteran insect pests through RNA interference. Nat Biotechnol. 2007;25:1322–6.

5. Mao Y-B, Cai W-J, Wang J-W, Hong G-J, Tao X-Y, Wang L-J, et al. Silencing a cotton bollworm P450 monooxygenase gene by plant-mediated RNAi impairs larval tolerance of gossypol. Nat Biotechnol. 2007;25:1307–13.

6. Jia R, Zhao H, Huang J, Kong H, Zhang Y, Guo J, et al. Use of RNAi technology to develop a PRSV-resistant transgenic papaya. Sci Rep. 2017;7:12636.

7. Zhang J, Khan SA, Hasse C, Ruf S, Heckel DG, Bock R. Pest control. Full crop protection from an insect pest by expression of long double-stranded RNAs in plastids. Science. 2015;347:991–4.

8. Guan R, Chu D, Han X, Miao X, Li H. Advances in the Development of Microbial Double-Stranded RNA Production Systems for Application of RNA Interference in Agricultural Pest Control. Front Bioeng Biotechnol [Internet]. 2021 [cited 2023 Jul 4];9. Available from: https://www.frontiersin.org/articles/10.3389/fbioe.2021.753790

9. Rodrigues TB, Mishra SK, Sridharan K, Barnes ER, Alyokhin A, Tuttle R, et al. First Sprayable Double-Stranded RNA-Based Biopesticide Product Targets Proteasome Subunit Beta Type-5 in Colorado Potato Beetle (Leptinotarsa decemlineata). Front Plant Sci. 2021;12:728652.

10. Arpaia S, Christiaens O, Giddings K, Jones H, Mezzetti B, Moronta-Barrios F, et al. Biosafety of GM Crop Plants Expressing dsRNA: Data Requirements and EU Regulatory Considerations. Front Plant Sci [Internet]. 2020 [cited 2023 Jul 4];11. Available from: https://www.frontiersin.org/articles/10.3389/fpls.2020.00940

11. Mehlhorn S, Ulrich J, Baden CU, Buer B, Maiwald F, Lueke B, et al. The mustard leaf beetle, Phaedon cochleariae, as a screening model for exogenous RNAi-based control of coleopteran pests. Pestic Biochem Physiol. 2021;176:104870.

12. Knorr E, Fishilevich E, Tenbusch L, Frey MLF, Rangasamy M, Billion A, et al. Gene silencing in Tribolium castaneum as a tool for the targeted identification of candidate RNAi targets in crop pests. Sci Rep. 2018;8:2061.

13. Ulrich J, Dao VA, Majumdar U, Schmitt-Engel C, Schwirz J, Schultheis D, et al. Large scale RNAi screen in Tribolium reveals novel target genes for pest control and the proteasome as prime target. BMC Genomics [Internet]. 2015 [cited 2016 Jun 15];16. Available from: http://www.biomedcentral.com/1471-2164/16/674

14. Christiaens O, Whyard S, Vélez AM, Smagghe G. Double-Stranded RNA Technology to Control Insect Pests: Current Status and Challenges. Front Plant Sci [Internet]. 2020 [cited 2023 Jul 4];11. Available from: https://www.frontiersin.org/articles/10.3389/fpls.2020.00451

15. Mehlhorn S, Hunnekuhl VS, Geibel S, Nauen R, Bucher G. Establishing RNAi for basic research and pest control and identification of the most efficient target genes for pest control: a brief guide. Front Zool. 2021;18:60.

16. Cooper AM, Silver K, Zhang J, Park Y, Zhu KY. Molecular mechanisms influencing efficiency of RNA interference in insects. Pest Manag Sci. 2019;75:18–28.

17. Bucher G, Scholten J, Klingler M. Parental RNAi in Tribolium (Coleoptera). Curr Biol. 2002;12:R85– 6.

18. Tomoyasu Y, Denell RE. Larval RNAi in Tribolium (Coleoptera) for analyzing adult development. Dev Genes Evol. 2004;214:575–8.

19. Tomoyasu Y, Miller SC, Tomita S, Schoppmeier M, Grossmann D, Bucher G. Exploring systemic RNA interference in insects: a genome-wide survey for RNAi genes in Tribolium. Genome Biol. 2008;9:R10.

20. Whangbo JS, Hunter CP. Environmental RNA interference. Trends Genet TIG. 2008;24:297–305.

21. Allen ML, Walker WB. Saliva of Lygus lineolaris digests double stranded ribonucleic acids. J Insect Physiol. 2012;58:391–6.

22. Garbutt JS, Bellés X, Richards EH, Reynolds SE. Persistence of double-stranded RNA in insect hemolymph as a potential determiner of RNA interference success: evidence from Manduca sexta and Blattella germanica. J Insect Physiol. 2013;59:171–8.

23. Prentice K, Christiaens O, Pertry I, Bailey A, Niblett C, Ghislain M, et al. RNAi-based gene silencing through dsRNA injection or ingestion against the African sweet potato weevil Cylas puncticollis (Coleoptera: Brentidae). Pest Manag Sci. 2017;73:44–52.

24. Prentice K, Smagghe G, Gheysen G, Christiaens O. Nuclease activity decreases the RNAi response in the sweetpotato weevil Cylas puncticollis. Insect Biochem Mol Biol. 2019;110:80–9.

25. Spit J, Philips A, Wynant N, Santos D, Plaetinck G, Vanden Broeck J. Knockdown of nuclease activity in the gut enhances RNAi efficiency in the Colorado potato beetle, Leptinotarsa decemlineata, but not in the desert locust, Schistocerca gregaria. Insect Biochem Mol Biol. 2017;81:103–16.

26. Wynant N, Santos D, Verdonck R, Spit J, Van Wielendaele P, Vanden Broeck J. Identification, functional characterization and phylogenetic analysis of double stranded RNA degrading enzymes present in the gut of the desert locust, Schistocerca gregaria. Insect Biochem Mol Biol. 2014;46:1–8.

27. Christiaens O, Tardajos MG, Martinez Reyna ZL, Dash M, Dubruel P, Smagghe G. Increased RNAi Efficacy in Spodoptera exigua via the Formulation of dsRNA With Guanylated Polymers. Front Physiol. 2018;9:316.

28. Lin Y-H, Huang J-H, Liu Y, Belles X, Lee H-J. Oral delivery of dsRNA lipoplexes to German cockroach protects dsRNA from degradation and induces RNAi response. Pest Manag Sci. 2017;73:960–6.

29. Zhang M, Koskie K, Ross JS, Kayser KJ, Caple MV. Enhancing glycoprotein sialylation by targeted gene silencing in mammalian cells. Biotechnol Bioeng. 2010;105:1094–105.

30. Schmitt-Engel C, Schultheis D, Schwirz J, Ströhlein N, Troelenberg N, Majumdar U, et al. The iBeetle large-scale RNAi screen reveals gene functions for insect development and physiology. Nat Commun. 2015;6:7822.

31. Bingsohn L, Knorr E, Billion A, Narva KE, Vilcinskas A. Knockdown of genes in the Toll pathway reveals new lethal RNA interference targets for insect pest control. Insect Mol Biol. 2017;26:92–102.

32. Nüsslein-Volhard C, Wieschaus E. Mutations affecting segment number and polarity in Drosophila. Nature. 1980;287:795–801.

33. Gonczy P, Echeverri C, Oegema K, Coulson A, Jones SJ, Copley RR, et al. Functional genomic analysis of cell division in C. elegans using RNAi of genes on chromosome III. Nature. 2000;408:331– 6.

34. Klingler M, Bucher G. The red flour beetle T. castaneum: elaborate genetic toolkit and unbiased large scale RNAi screening to study insect biology and evolution. EvoDevo. 2022;13:14.

35. Tomoyasu Y, Denell RE. Larval RNAi in Tribolium (Coleoptera) for analyzing adult development. Dev Genes Evol. 2004;214:575–8.

36. Hakeemi MS, Ansari S, Teuscher M, Weißkopf M, Großmann D, Kessel T, et al. Screens in fly and beetle reveal vastly divergent gene sets required for developmental processes. BMC Biol. 2022;20:38.

37. Götz S, García-Gómez JM, Terol J, Williams TD, Nagaraj SH, Nueda MJ, et al. High-throughput functional annotation and data mining with the Blast2GO suite. Nucleic Acids Res. 2008;36:3420–35.

38. Kanehisa M, Goto S. KEGG: Kyoto Encyclopedia of Genes and Genomes. Nucleic Acids Res. 2000;28:27–30.

39. dos Santos G, Schroeder AJ, Goodman JL, Strelets VB, Crosby MA, Thurmond J, et al. FlyBase: introduction of the Drosophila melanogaster Release 6 reference genome assembly and large-scale migration of genome annotations. Nucleic Acids Res. 2015;43:D690–7.

40. Supek F, Bošnjak M, Škunca N, Šmuc T. REVIGO summarizes and visualizes long lists of gene ontology terms. PloS One. 2011;6:e21800.

41. Abd El Halim HM, Alshukri BMH, Ahmad MS, Nakasu EYT, Awwad MH, Salama EM, et al. RNAi-mediated knockdown of the voltage gated sodium ion channel TcNav causes mortality in Tribolium castaneum. Sci Rep. 2016;6:29301.

42. Cao M, Gatehouse JA, Fitches EC. A Systematic Study of RNAi Effects and dsRNA Stability in Tribolium castaneum and Acyrthosiphon pisum, Following Injection and Ingestion of Analogous dsRNAs. Int J Mol Sci. 2018;19:1079.

43. Peng Y, Wang K, Chen J, Wang J, Zhang H, Ze L, et al. Identification of a double-stranded RNA-degrading nuclease influencing both ingestion and injection RNA interference efficiency in the red flour beetle Tribolium castaneum. Insect Biochem Mol Biol. 2020;125:103440.

44. Whyard S, Singh AD, Wong S. Ingested double-stranded RNAs can act as species-specific insecticides. Insect Biochem Mol Biol. 2009;39:824–32.

45. Zhu F, Xu J, Palli R, Ferguson J, Palli SR. Ingested RNA interference for managing the populations of the Colorado potato beetle, Leptinotarsa decemlineata. Pest Manag Sci. 2011;67:175–82.

46. Herndon N, Shelton J, Gerischer L, Ioannidis P, Ninova M, Dönitz J, et al. Enhanced genome assembly and a new official gene set for Tribolium castaneum. BMC Genomics. 2020;21:47.

47. Hu X, Richtman NM, Zhao J-Z, Duncan KE, Niu X, Procyk LA, et al. Discovery of midgut genes for the RNA interference control of corn rootworm. Sci Rep. 2016;6:30542.

48. Rodrigues TB, Duan JJ, Palli SR, Rieske LK. Identification of highly effective target genes for RNAi-mediated control of emerald ash borer, Agrilus planipennis. Sci Rep. 2018;8:5020.

49. Pridgeon JW, Zhao L, Becnel JJ, Strickman DA, Clark GG, Linthicum KJ. Topically applied AaeIAP1 double-stranded RNA kills female adults of Aedes aegypti. J Med Entomol. 2008;45:414–20.

50. Tian H, Peng H, Yao Q, Chen H, Xie Q, Tang B, et al. Developmental Control of a Lepidopteran Pest Spodoptera exigua by Ingestion of Bacteria Expressing dsRNA of a Non-Midgut Gene. PLoS ONE. 2009;4:e6225.

51. Kumar M, Gupta GP, Rajam MV. Silencing of acetylcholinesterase gene of Helicoverpa armigera by siRNA affects larval growth and its life cycle. J Insect Physiol. 2009;55:273–8.

52. Yu R, Xu X, Liang Y, Tian H, Pan Z, Jin S, et al. The Insect Ecdysone Receptor is a Good Potential Target for RNAi-based Pest Control. Int J Biol Sci. 2014;10:1171–80.

53. Beder T, Aromolaran O, Dönitz J, Tapanelli S, Adedeji EO, Adebiyi E, et al. Identifying essential genes across eukaryotes by machine learning. NAR Genomics Bioinforma. 2021;3:lqab110.

54. Boaventura D, Buer B, Hamaekers N, Maiwald F, Nauen R. Toxicological and molecular profiling of insecticide resistance in a Brazilian strain of fall armyworm resistant to Bt Cry1 proteins. Pest Manag Sci. 2021;77:3713–26.

55. Conesa A, Gotz S, Garcia-Gomez JM, Terol J, Talon M, Robles M. Blast2GO: a universal tool for annotation, visualization and analysis in functional genomics research. Bioinformatics. 2005;21:3674–6.

56. Quevillon E, Silventoinen V, Pillai S, Harte N, Mulder N, Apweiler R, et al. InterProScan: protein domains identifier. Nucleic Acids Res. 2005;33:W116–120.

57. Mahram A, Herbordt MC. NCBI BLASTP on High-Performance Reconfigurable Computing Systems. ACM Trans Reconfigurable Technol Syst. 2015;7:33:1–33:20.

58. Young MD, Wakefield MJ, Smyth GK, Oshlack A. Gene ontology analysis for RNA-seq: accounting for selection bias. Genome Biol. 2010;11:R14.

