## Supporting figures and text for "Superior target genes and pathways for RNAi mediated pest control revealed by genome wide analysis in the red flour beetle *Tribolium castaneum*"

### Supporting information 1 - Supporting figures

#### Supporting figure 1: Schedule and procedure for the high-throughput primary screen.

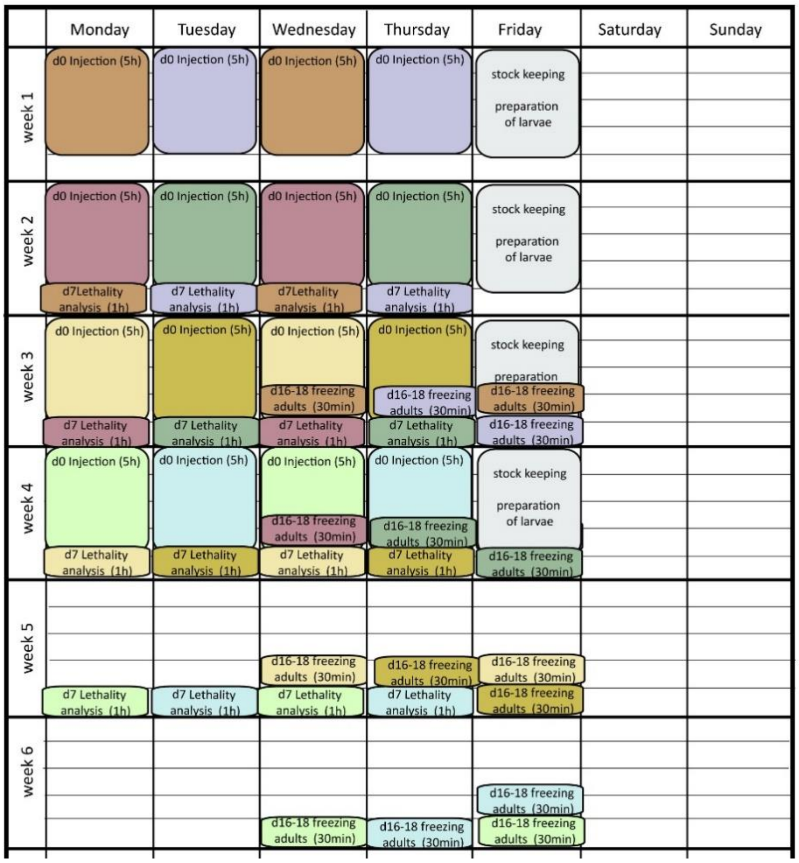


Schedule depicting the different processing steps - one set of parallel experiments is shown in the same color. In this depiction, 16 such parallel experiments are shown (i.e. 4 per week).

On each injection day (d0), dsRNAs targeting 39 different genes plus one buffer control were injected into 10 larvae of L6 each (L5 used only in case of need). Injection was done four days a week (Monday-Thursday) while the fifth day was used for stock keeping and collection of larvae for the following week's injections. The first injection of each day was a buffer control, which served as negative control and was at the same time needed as warm-up for the injection procedure (We had previously observed a generally increased level of background lethality in the first injection of a day).

Seven days later (d7) the survival was scored for the first time while 16-18 days later survival was checked a second time (data not shown) and the pupae/adults were frozen for later morphological analyses.

The schedules were interleaved such that week 5 of the first round of experiments is at the same time week 1 of the following repetition of the schedule (not shown). This procedure was performed by one person for approximately two years showing that the workload of screening 156 genes per week (plus controls) was sustainable. We would consider it challenging to increase the throughput if the screener had to do it over extended time considering European worker protection standards.

#### Supporting Figure 2: Lethality distribution of the primary screen as basis for the selection for the validation screen

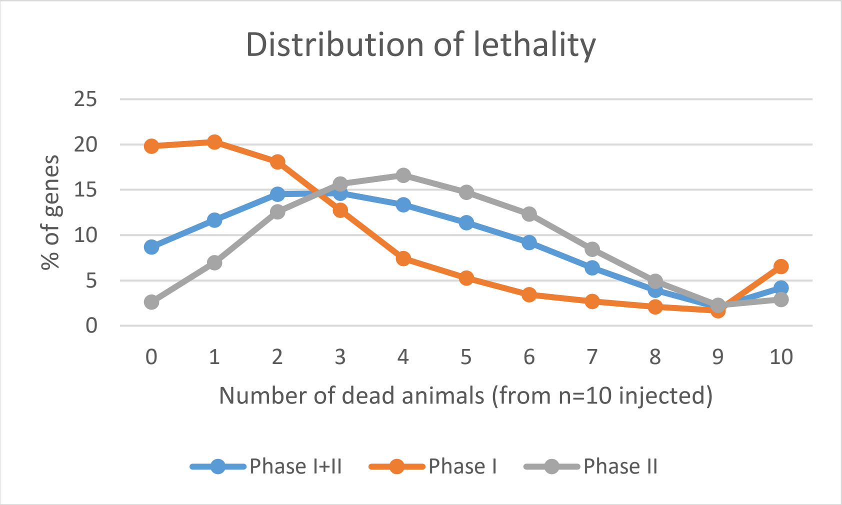


In order to determine the cutoff for selecting target genes for the validation screen, we used the distribution of dead animals per experiment for the earliest available lethality data for phase I (11 days post injection, orange line), phase II (7 days post injection grey line) and combined those datasets (blue line). We expected a distribution of technical or background lethality in addition to the gene specific RNAi effects. Indeed, we found that the lethality distribution of all datasets approached a minimum at 90 % dead animals (9 out of 10 injected) while the value for 100 % lethality was higher again. This increase of the 100 % value is best explained by RNAi induced lethality in those datasets. The number of experiments with a 90 % value was not clearly increased and therefore likely contained a portion of false positive datasets, i.e. data where technical lethality had contributed to the signal. Hence, for the validation screen we considered all genes showing 100 % lethality as candidates but still included some genes with 90 % lethality in order not to miss good candidates.

### Supporting Figure 3: Lethality distribution of the validation screen

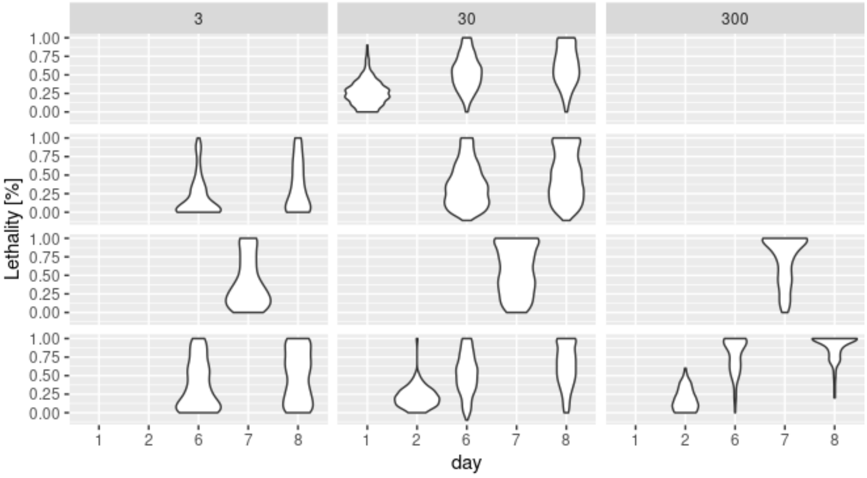


The validation screen (total n=843 genes) was performed in different phases by different people and under changing organizational constraints. In the first and second part, all three concentrations were used (bottom two rows). As the higher concentration seemed to reflect the results of the primary screen, that concentration was skipped in the subsequent phases of the validation screen (two top rows). The last part of the validation screen was performed only for the 30 ng/ul concentration (top row; n=400), which had shown to relieably separate very good from less efficacious target genes.

#### Supporting Figure 4: Revigo network of GO terms "biological process"

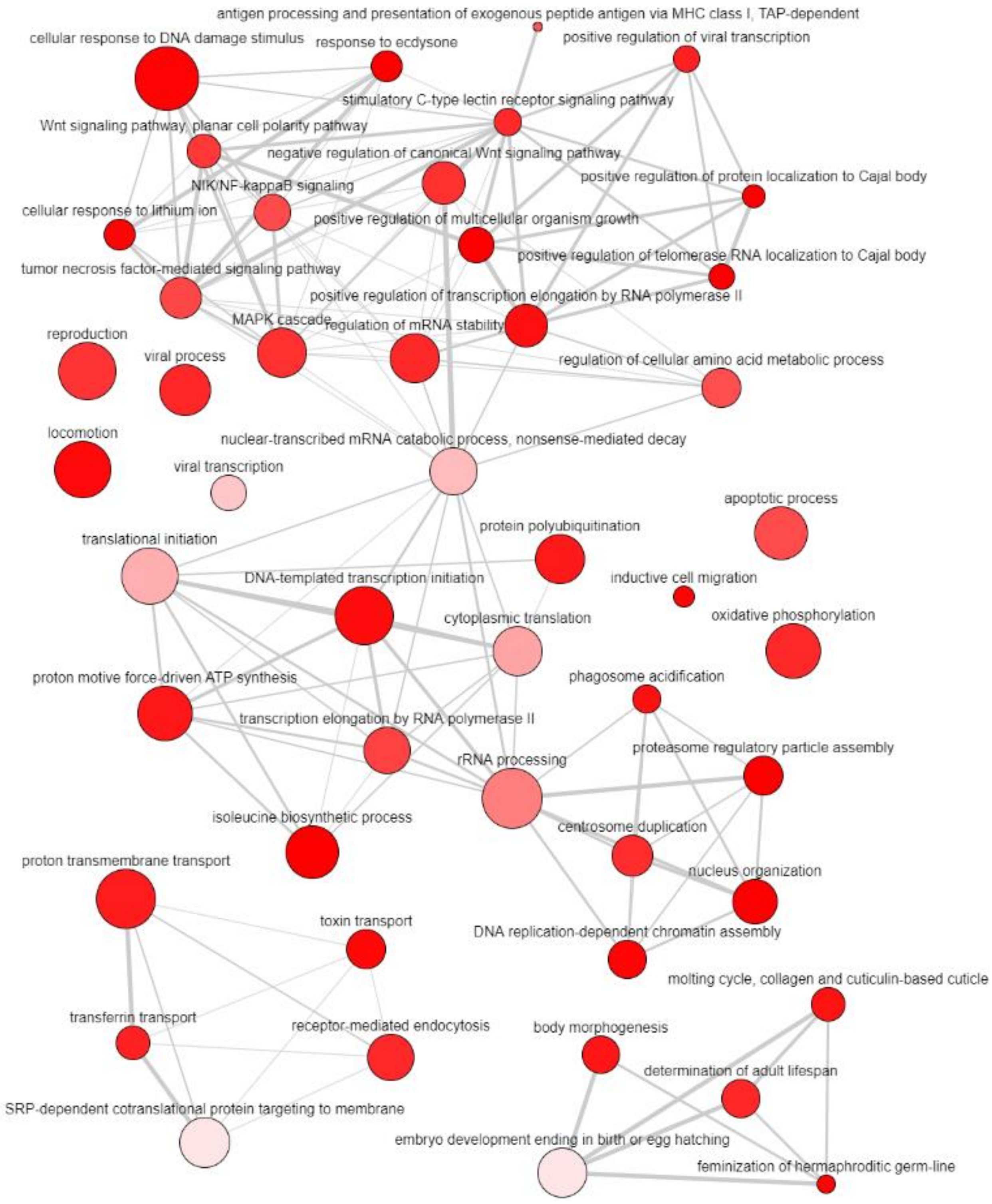


The color of the bubble corresponds to the p-value as given in the Supplementary Table 1: More intense colors reflects higher confidence. The size of the bubble corresponds to the LogSize p-value for the GO Term.

#### Supporting Figure 5 Revigo network of GO terms “cellular component”

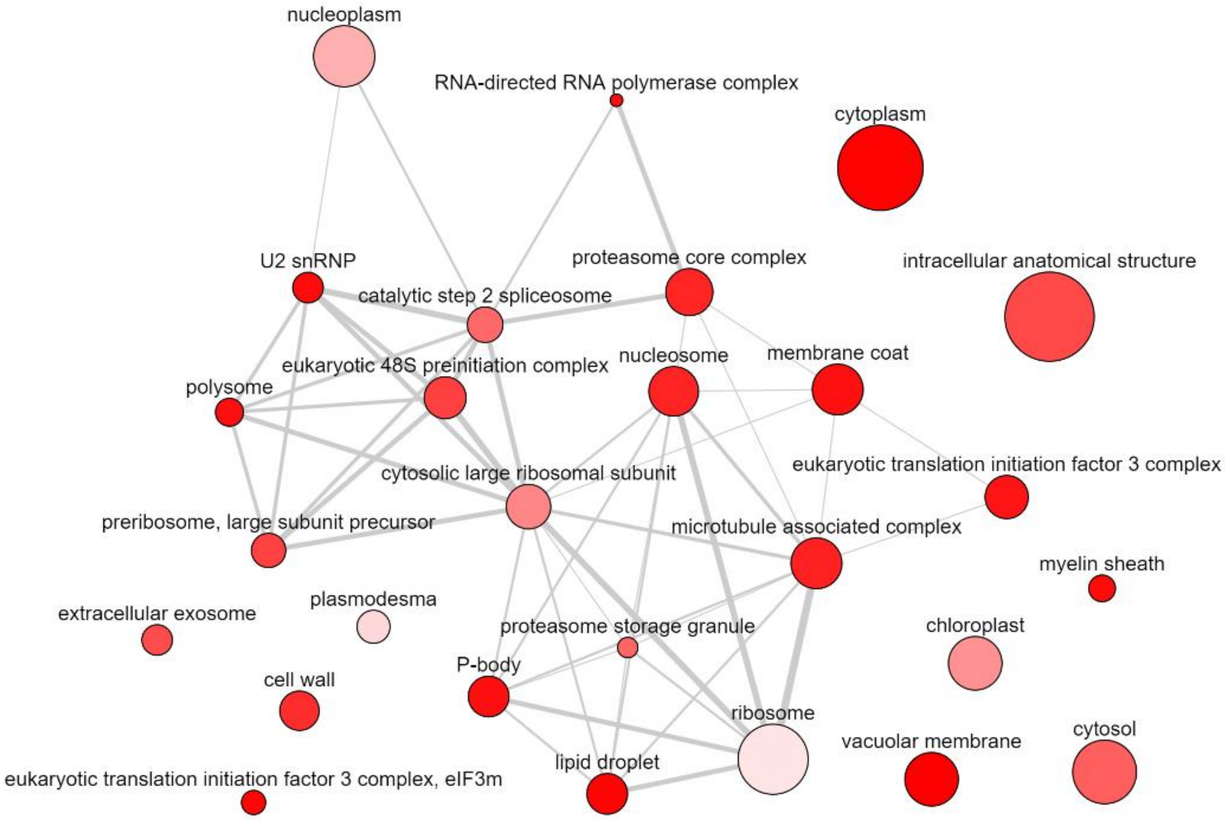


The color of the bubble corresponds to the p-value as given in the Supplementary Table 1: More intense colors reflects higher confidence. The size of the bubble corresponds to the LogSize p-value for the GO Term.

#### Supporting Figure 6: Revigo network of GO terms "molecular function"

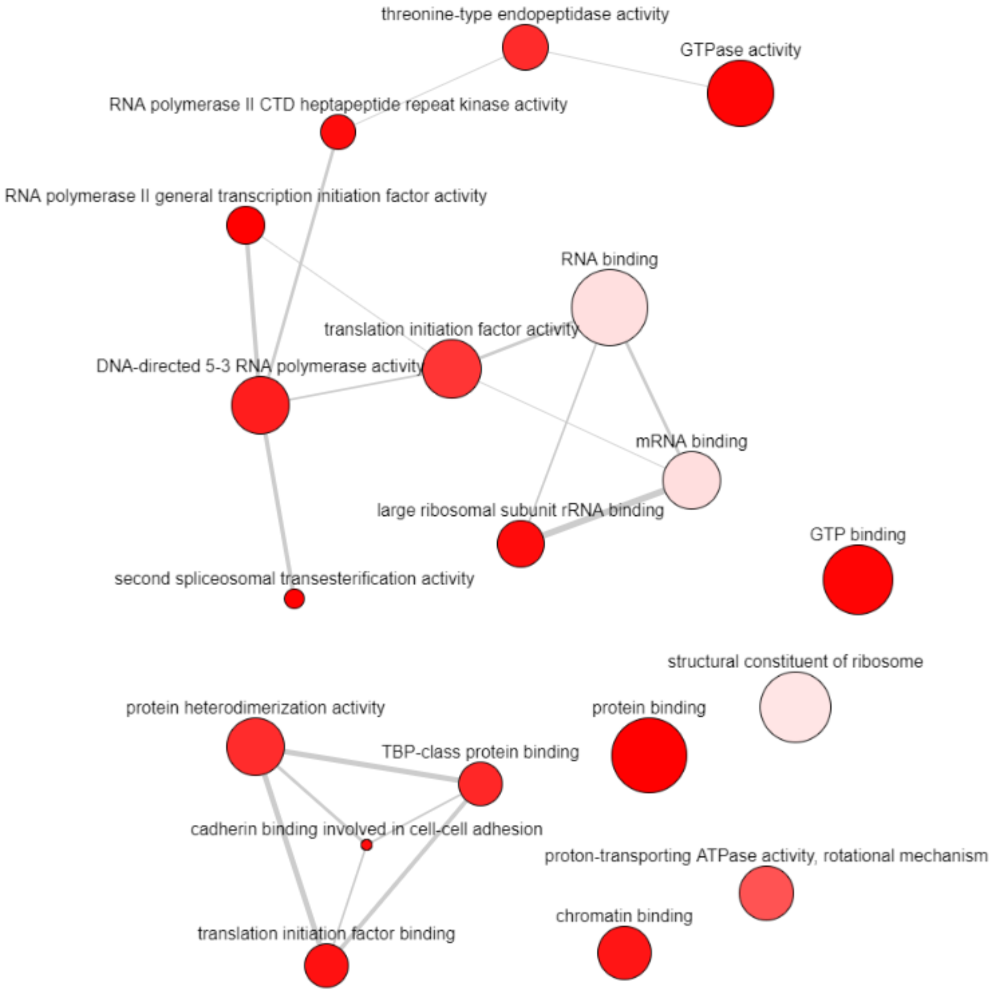


The color of the bubble corresponds to the p-value as given in the Supplementary Table 1: More intense colors reflects higher confidence. The size of the bubble corresponds to the LogSize p-value for the GO Term.
